# An emerging role of *Prevotella histicola* on estrogen-deficiency induced bone loss through the gut microbiota-bone axis

**DOI:** 10.1101/2020.06.03.133082

**Authors:** Zhongxiang Wang, Kai Chen, Congcong Wu, Junhao Chen, Hao Pan, Yangbo Liu, Peng Wu, Jiandong Yuan, Junzhe Lang, Juanjuan Du, Jiake Xu, Keke Jin, Lei Chen

**Affiliations:** Department of Orthopaedics, The First Affiliated Hospital of Wenzhou Medical University, Wenzhou 325000, China; School of Biomedical Sciences, The University of Western Australia, Perth, WA 6009, Australia; Department of Pathophysiology, Wenzhou Medical University, Wenzhou 325000, China; Cardiac Regeneration Research Institute, Wenzhou Medical University, Wenzhou 325000, China; Institute of ischemia/reperfusion injury, Wenzhou Medical University, Wenzhou 325000, China; Nervous Institute in Basic College, Wenzhou Medical University, Wenzhou 325035, China

**Keywords:** Gut microbiota, Postmenopausal osteoporosis, Osteoclastogenic cytokines, Gut permeability, prevotella histicola

## Abstract

Gut microbiota (GM)-bone axis has emerged as a crucial mediator in the maintenance of bone homeostasis. Estrogen-deficiency induced bone loss is closely associated with an altered GM. However, the underlying mechanisms remain not fully understood. To this end, feces samples collected from the postmenopausal patients with osteoporosis (PMO) and with normal bone mass (PMN) were examined by 16s rRNA gene sequencing. We found that GM in PMO group was featured by a significant decreased proportion of genus *Prevotella* in comparison with that in the PMN group. Next, *Prevotella histicola* (*P.histicola*), a typical specie of *Prevotella*, was orally given to the mice following the ovariectomy (OVX) procedure and it significantly prevented OVX induced bone loss. Mechanistically, the protective effects of *P.histicola* on bone mass were found to be associated with the inhibition of osteoclastic resorption by attenuating osteoclastogenic cytokines expression and ameliorating gut permeability. Thus, *P.histicola* may prevent estrogen-deficiency induced bone loss through the GM-bone axis.

## Introduction

Osteoporosis is known as a systemic skeletal disease characterized by low bone mass, which eventually leads to an increased bone fragility and becomes susceptible to fractures (Leslie & Morin, 2014). In USA, it was estimated that 40% of women over the of age 50 will suffer an osteoporotic fracture which is closely associated with high morbidity and mortality (Cheng, Wentworth et al., 2020). Despite current treatments on promoting bone health and reducing fractures, the number of postmenopausal osteoporosis (PMO) patients is still rising worldwide due to the aging population. However, unwanted medication side effects may largely limit their safety for long-term use (Cheng et al., 2020). Novel effective approaches with fewer side effects are urgently needed.

Excessive osteoclast formation and resorption are considered as the key pathological changes in estrogen-deficiency induced osteoporosis (Feng, Liu et al., 2014). Receptor activator of nuclear factor kappa-Β (NF-*κ*B) ligand (RANKL), majorly sourced from osteoblast lineage cells, is an indispensable factor involved in osteoclast formation by binding to RANK - its receptor on the surface of osteoclast precursors (Yasuda, Shima et al., 1998). This effect is opposed by the decoy receptor for RANKL - osteoprotegerin (OPG) which is derived from marrow stromal cells (MSCs) and osteoblasts (Eghbali-Fatourechi, Khosla et al., 2003). Estrogen efficiently reduces the source of RANKL but increases the production of OPG, thereby mediating osteoclast formation and function (Liu, Zhang et al., 2003). Besides, estrogen was also demonstrated to modulate the production of a series of osteoclastogenic cytokines including tumor necrosis factor-*α* (TNF*α*), interleukin (IL)-1β and IL-6 (Faienza, Ventura et al., 2013) which act as pro-resorptive factors by enhancing RANKL expression in osteoblast lineage cells (Boyle, Simonet et al., 2003). Furthermore, estrogen-deficiency leads to compromised gut integrity and result in microbiota-associated cytokines passing through the gap junctions. (Li, Chassaing et al., 2016b).

Interestingly, it was widely accepted that osteoporosis does not develop in every postmenopausal woman. In our study, 42.86% (18/42) of postmenopausal women presented the normal BMD, which means some indispensable factors may promote the bone loss in addition to estrogen deficiency. Gut microbiota (GM), composed of trillions of bacteria in gastrointestinal tract (Qin, Li et al., 2010), is considered as a crucial determinant for human health (Ley, Hamady et al., 2008). Accumulating evidence suggested that altered GM compositions largely contribute to various metabolic disorders including inflammatory bowel disease (IBD), obesity, diabetes and osteoporosis (Huang, Inoue et al., 2015, Kang, Jeraldo et al., 2014, Spor, Koren et al., 2011, Wang, Wang et al., 2017). GM-bone axis was previously found to mediate the estrogen-deficiency induced osteoporosis. Compared with normal mice, germ-free (GF) mice showed less bone loss following estrogen-deficiency due to the reduction of osteoclastogenic cytokines (Sjogren, Engdahl et al., 2012). In addition, probiotics treatment could reduce the expression of osteoclastogenic cytokines and increase the expression of OPG in bone, protecting mice from ovariectomy (OVX)-induced bone loss (Ohlsson, Engdahl et al., 2014). Hence, GM may serve as a prerequisite for estrogen-deficiency induced osteoporosis and therapeutic strategies based on GM-bone axis may be promising in osteoporosis treatment. However, it still remains largely unclear how GM profiles change in PMO patients and to what extent can the supplement of a significant microbiome ameliorate the bone loss.

Given the close relationship between GM and PMO, we hypothesized that the compositions of GM in PMO patients may be largely altered which increased the susceptibility to osteoporosis. To better characterize the GM compositions in PMO patients and advance the understanding of the GM-bone axis, we analyzed the diversity and richness of GM from the PMO patients and postmenopausal women with normal bone mass (PMN). We found that GM in PMO group was featured by a significantly decreased proportion of genus *Prevotella* compared with that in PMN group. Furthermore, *Prevotella histicola* (*P.histicola*), a specie of *Prevotella*, was used to orally treat the mice after the OVX procedure, and we found it was able to significantly prevent OVX induced bone loss. Mechanistically, *P.histicola* inhibited osteoclast activity by ameliorating gut permeability and attenuating osteoclastogenic cytokines. Thus, our study highlighted a protective role of *P.histicola* on estrogen-deficiency induced bone loss through GM-bone axis.

## Results

### Diversity analysis of GM derived from PMN and PMO

Fecal samples were collected from PMO group (n=24) and PMN group (n=18), and then proceeded to 16S rRNA sequencing and analysis (Fig. 1a). We also have tested the BMD and some serum indexes related to bone remodeling. Characteristics and clinical data of participants are provided in Table 1. Alter filtering, Illumina MiSeq sequencing yielded a total of 1, 270, 876 high-quality reads from 42 samples with an average of 30, 259 reads in every sample, and 962 operational taxonomic units (OTUs) were identified. Alpha analyses were performed to compare the differences of diversity and richness between PMO and PMN groups. It was indicated that the richness of GM in PMN group was significantly higher than that in the PMO group, as evidenced by Ace and Chao index; and Shannon and Simpson index revealed little difference in diversity between these two groups (Fig. 1b). These data suggested that the development of osteoporosis may be mainly related with an altered richness of GM rather than the diversity.

**Table 1.** Characteristics of participants

|  | **Normal Control** | **Osteoporosis** | **P** |
| --- | --- | --- | --- |
| Age, y | 61.51±4.13 | 60.12±4.59 | >0.05 |
| Height, cm | 157.39±3.93 | 154.83±5.95 | >0.05 |
| Weight, kg | 56.44±5.46 | 52.06±9.74 | >0.05 |
| BMI, kg/m<sup>2</sup> | 22.76±1.69 | 21.80±4.32 | >0.05 |
| BMD of lumbar vertebrae, T-score | 0.20±1.03 | -3.13±0.78 | <0.001 |
| BMD of femoral neck, T-score | -0.31±0.62 | -2.45±0.78 | <0.001 |
| PINP of serum, ng/mL | 52.46±26.12 | 64.41±34.43 | >0.05 |
| β-CTX of serum, ng/mL | 0.54±0.23 | 0.60±0.25 | >0.05 |
| Calcium of serum, mmol/L | 2.18±0.09 | 2.18±0.13 | >0.05 |
| Phosphorus of serum, mmol/L | 1.03±1.74 | 7.60±31.6 | >0.05 |

**Figure 1.**
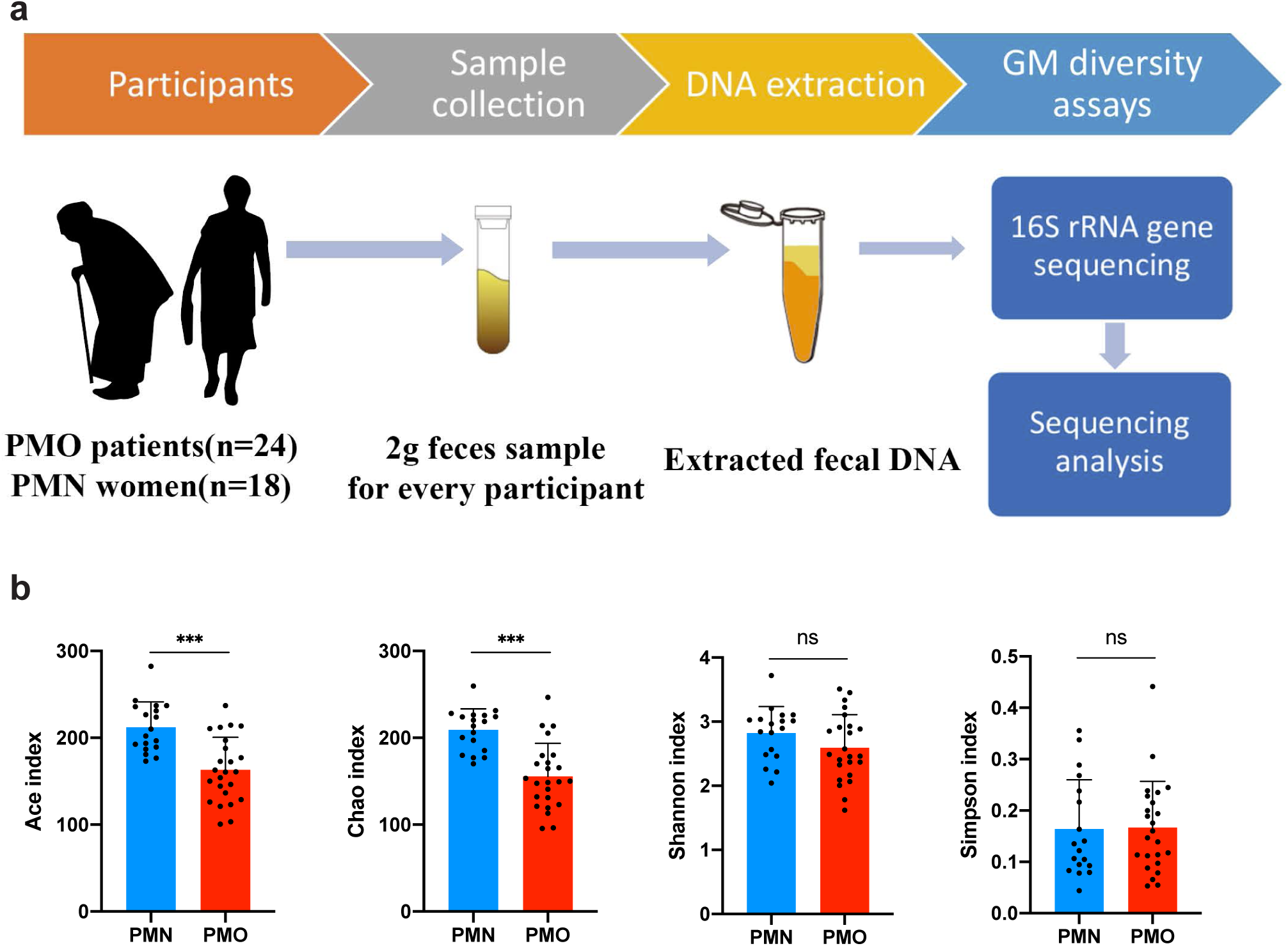
An altered richness in the gut microbiota (GM) of the postmenopausal osteoporosis population. **a** The flow chart depicting the participants selection, fecal microbiota DNA preparation and GM 16S rRNA gene sequencing. **b** Alpha diversity analysis of GM between postmenopausal women with osteoporosis (PMO) (n=24) and normal bone mass, as normal control(NC) (n=18) showing that the richness of PMO group is significantly decreased as indicated by Ace and Chao indexes, while the abundance remain insignificantly different between these two groups as indicated by Shannon and Simpson indexes. (**** p<0.0001)

### Identification of representative bacterial taxa between two groups

Linear discriminant analysis of Effect Size (LEfSe) was used to compare the different biomarkers between groups. We found that 21 taxa were under-represented and 9 were over-represented in PMO patients (Fig. 2a and b). At the family level, GM of PMN group was enriched in *Prevotellaceae, Acidaminococcaceae, Coriobacteriaceae* and *Hydrogenophilaceae*, whereas GM of PMO group were prominent with *Bacteroidaceae, Hyphomicrobiaceae* and *Pasteurellaceaae*. All of these families of bacteria were key phylotypes involved in the segregation of GM in PMN and PMO patients following the LEfSe analysis. Of note, the richness of *Prevotellaceae* remained the lowest in the PMO group as compared with that in PMN group, which indicated that it may play a protective role on the estrogen-deficiency induced bone loss.

**Figure 2.**
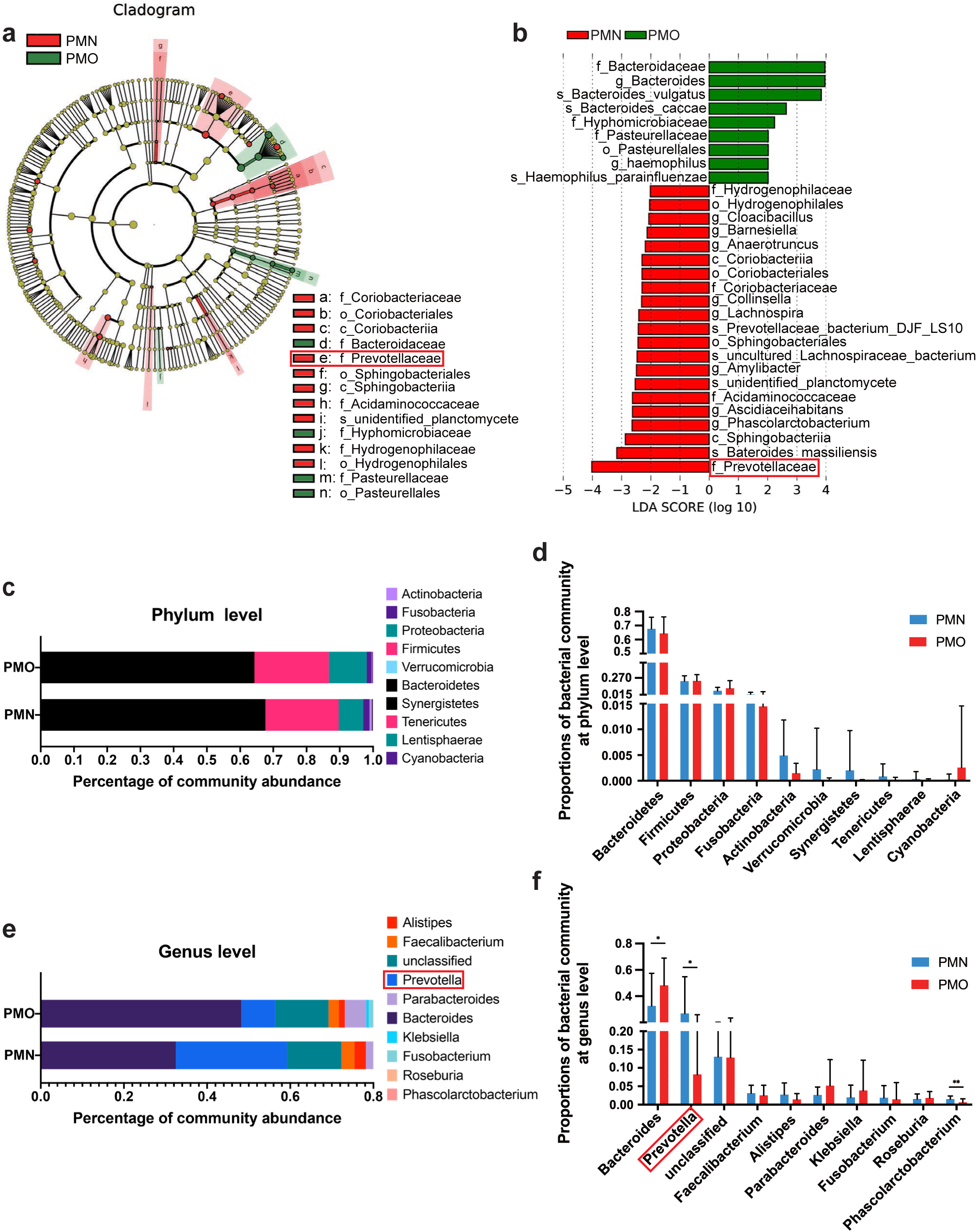
A decreased proportion of genus *Prevotella* in the patients with postmenopausal osteoporosis (PMO). **a** Taxonomic representation of statistically and biologically consistent differences between two groups. Differences are represented by the color of the most abundant class (red indicating group N, green group O and yellow non-significant). The diameter of each circle’s diameter is proportional to the taxon’s abundance. **b** Histogram of the LDA scores for abundance differences at the level of class, order, family, genus and species. Cladogram was calculated by LefSe, and displayed according to effect size. **c** Gut bacterial community compositional at phylum level. **d** Significant difference of top 10 bacterial community richness at phylum level. **e** Gut bacterial community compositional at genus level. f Significant difference of top 10 bacterial community richness at genus level. (* p<0.05, ** p<0.01)

### A decreased proportion of genus *Prevotella* in the GM of PMO group

To further compare the differences of microbial compositions between the PMN and PMO groups, we next analyzed the bacterial communities by performing taxonomic assignment of all reads. The results showed that, at the phylum level, the two groups were both majorly composed of *Bacteroides, Firmicutes, Proteobacteria, Fusobacteria, Actinobacteria* and *Verrucomicrobia*, and the compositions were insignificantly different (Fig. 2c and d). However, the proportions of bacterial community between the two groups are different at the genus level. A total of 53 genus were detected with a ratio above 0.1%, among which the genus of *Bacteroides* and *Prevotella* predominantly contributed more than 60%. The GM of PMO women displayed an obviously decrease in *Prevotella* and an increase in *Bacteroides* compared to PMN patients. It was noticeable that the richness of *Prevotella* of GM in PMN women was more than three times than that of PMO patients. These data indicated that the GM compositions have changed in PMO patients, and genus *Prevotella* is supposed to play a protective role.

### *P.histicola* prevents OVX-induced bone loss

To identify the potential effects of *Prevotella* on estrogen-deficiency induced bone loss and better understand the underlying mechanisms, *P.histicola* - a typical species of *Prevotella* - was orally administrated in the OVX mice and the samples including gut, serum, and bone were collected for further analysis (Fig. 3a). *P.histicola* has been widely shown to alleviate the arthritis and demyelinating disease in mice via suppressing inflammation in previous studies (Mangalam, Shahi et al., 2017, Marietta, Murray et al., 2016). To characterize the bone mass of the femur in different groups, micro-CT scanning and Van Gieson (VG) staining of the femur were performed and the results showed the bone volume was largely reduced following OVX procedures while the use of *P.histicola* effectively prevented the OVX mice from bone loss (Fig. 3b-g). These changes were well evidenced by the quantitative analysis of bone parameters including bone volume per tissue volume (BV/TV), trabecular number (Tb.N), trabecular space (Tb.Sp) (Fig. 3c-e). However, the trabecular thickness (Tb.Th) remained unchanged in our study (Fig. 3f). These data indicated that the supplement of *P.histicola* is able to prevent estrogen-deficiency induced bone loss.

**Figure 3.**
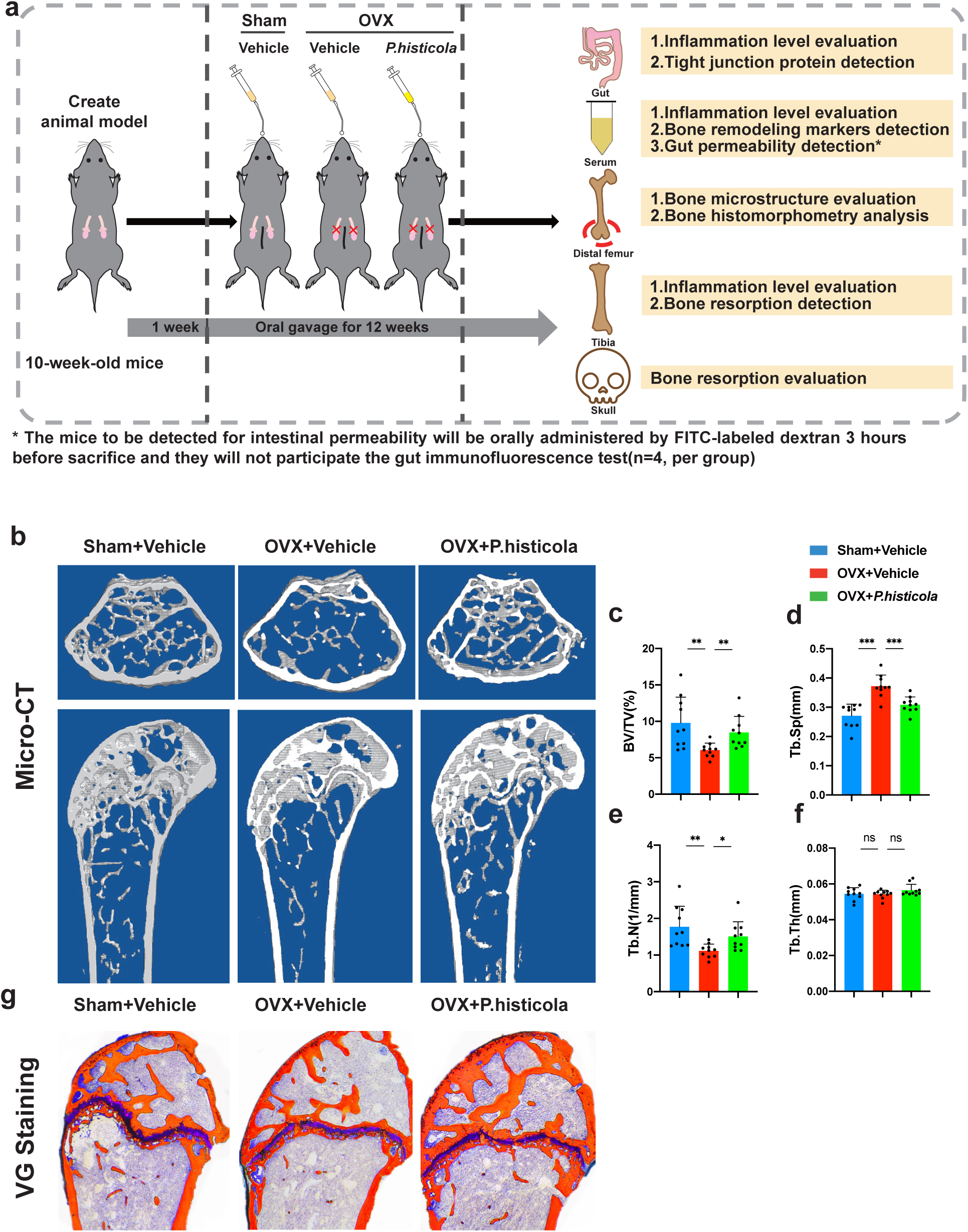
*Prevotella histicola* (*P.histicola*) prevents ovariectomy (OVX) induced bone loss. **a** Schematic illustration of the establishment of OVX mouse model and the experimental design to evaluate *P.histicola*’s effects. **b** Representative Micro-CT images showing the bone loss following OVX procedure and the supplement of *P.histicola* greatly prevent these changes (n=10 per group). **c-f** Quantitative analyses of parameters regarding bone microstructure, including BV/TV, Tb.Sp, Tb.N, Tb.Th (n=10 per group). **g**. Representative histological images of distal femurs stained by Van Gieson (n=3 per group). *** p<0.001, ** p<0.01, * p<0.05.BV/TV, bone volume per tissue volume; Tb.Sp, trabecular separation; Tb.N, trabecular number; Tb.Th, trabecular thickness; ns, non-significant.

### *P.histicola* reduces osteoclast activity in OVX mice through modulating RANKL/RANK/OPG pathway

Previous studies have shown that estrogen-deficiency induced bone loss was mainly caused by excessive osteoclast formation and resorption (Feng et al., 2014). To examine whether treatment with *P.histicola* could affect osteoclast activity, the bone sections of the distal femurs as well as the skull were stained using TRAP solution. The results showed the OVX procedure caused an increased activity of TRAP-positive osteoclasts, and *P.histicola* treatment efficiently inhibited the osteoclast activity (Fig. 4a). Quantitative analysis confirmed that the osteoclast number and surface area in the OVX mice were reduced by *P.histicola* treatment (Fig. 4b and c). Furthermore, serum biomarkers including N-terminal propeptide of type I procollagen (PINP) and C-terminal telopeptide of type I collagen (CTX-1) were detected to evaluate the bone remodeling. PINP, an osteoblast-related protein, showed no significance among three groups (Fig. 4d) while the osteoclastic resorption marker CTX-1 showed the consistent trend with the osteoclast activity in histological analysis (Fig. 4e), suggesting *P.histicola* may prevent osteoporosis by targeting osteoclasts rather than osteoblasts. RANKL/RANK/OPG axis is essential for the osteoclast formation and function. RANKL is indispensable for the osteoclastogenesis by binding to RANK, while OPG prevents the interaction of RANKL with RANK (Eghbali-Fatourechi et al., 2003, Yasuda et al., 1998). The OVX mice showed increased mRNA expressions of RANK as well as the RANKL/OPG ratio, which can be reversed by *P.histicola* treatment (Fig. 4f and g). These data indicated that *P.histicola* suppresses osteoclast activity by targeting RANKL/RANK/OPG pathway.

**Figure 4.**
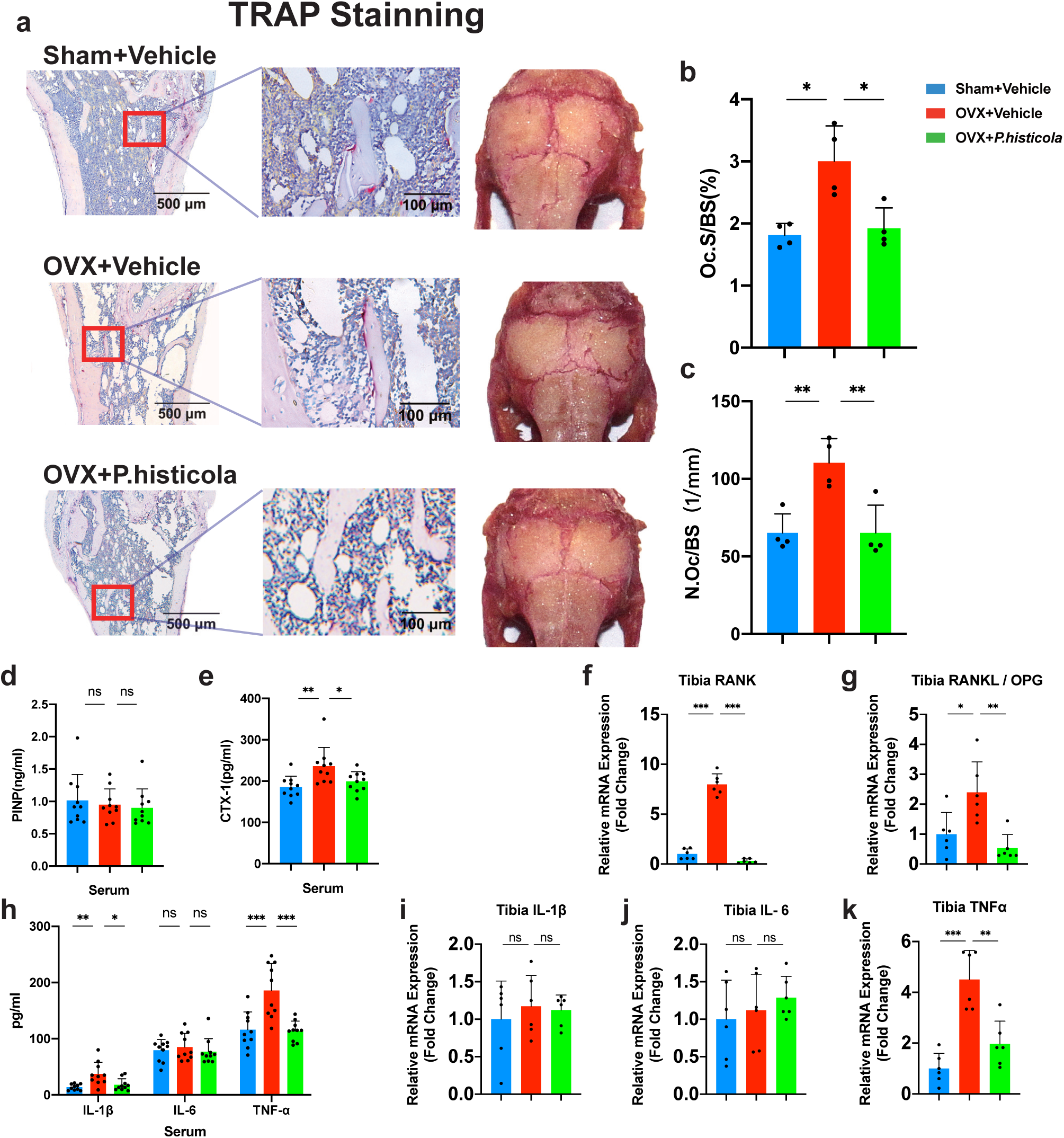
*Prevotella histicola* (*P.histicola*) inhibits osteoclast formation by suppressing the inflammation level in ovariectomy (OVX) mice. **a** The TRAP stainning showing that *P.histicola* inhibit the osteoclast formation on the bone surface in the OVX group (n=3 per group). **b-c** Quantitative analyses of osteoclast parameters including N.Oc/BS and Oc.S/B (n=4 per group). **d-e** Peripheral blood level of bone formation maker P1NP and bone resorption marker CTX-1(n=10 per group). **f-h** Levels of pro-resorptive cytokines RANK, RANKL and OPG expression in tibia among three groups(n=6 per group). **i** The levels of osteoclastogenesis related inflammatory cytokines in peripheral blood(n=10 per group). **j-l** Levels of osteoclastogenic cytokines IL-1β, IL-6 and TNFα expression in tibia among three groups(n=6 per group). *** p<0.001, ** p<0.01, * p<0.05. TRAP, tartrate-resistant acid phosphatase; N.Oc/BS, osteoclast number/bone surface; Oc.S/BS, osteoclast surface/bone surface; IL-1β, interluekin-1β; IL-6, interluekin-6; TNFα, tumor necrosis factor α. RANK, receptor activator of nuclear factor κB; RANKL, receptor activator of nuclear factor κB ligand; OPG, osteoprotegerin; P1NP, N-terminal propeptide of type 1 procollagen; CTX-1, cross-linked C-terminal telopeptide of type I collagen; ns, non-significant; P.histicola, Prevotella histicola.

### *P.histicola* inhibits the osteoclastogenic cytokines in serum and bone

It was reported that many inflammatory cytokines were able to promote osteoclastogenesis by enhancing RANKL/RANK/OPG pathway(Hofbauer, Lacey et al., 1999, Kwan Tat, Padrines et al., 2004, Pacifici, 2012, Sherman, Weber et al., 1990), particularly IL-1β, IL-6 and TNF*α* which were also known as osteoclastogenic cytokines (Ammann, Rizzoli et al., 1997, Kimble, Srivastava et al., 1996, Kimble, Vannice et al., 1994, Manolagas & Jilka, 1995, Tanaka, Takahashi et al., 1993). To determine how the treatment of *P.histicola* affects these cytokines, we assessed the serum level of IL-1β, IL-6 and TNF*α*. The results showed that ovariectomy augmented serum TNF*α* and IL-1β levels compared to the sham group, and *P.histicola* treatment could defy the increase of TNF*α* and IL-1β. The serum IL-6 level remained insignificantly changed among three groups (Fig. 4h). This was further supported by the qPCR results which examined these cytokines in the bone tissues at gene level (Fig. 4i, j and k). These results suggested that *P.histicola* suppresses the osteoclastogenesis by reducing the systemic levels of osteoclastogenic cytokines.

### *P.histicola* alters the expression of osteoclastogenic cytokines including IL-1β and TNFα in gut site -specifically

To investigate whether the changes of systemic osteoclastogenic cytokines in serum and bone resulted from the alterations of the gut, we next evaluated mRNA transcripts of IL-1β, IL-6 and TNF*α* from different parts of the gut including duodenum, jejunum, ileum, and colon. The mRNA expressions of these cytokines showed no significant change in the duodenum (Fig. 5a). However, in jejunum and ileum, mice following OVX procedure showed an increased IL-1β expression which was suppressed by *P.histicola* (Fig. 5 b and c). The expression of TNF*α*, a cytokine which can directly enhance osteoclastogenesis, was attenuated by *P.histicola* in both ileum and colon (Fig. 5 c and d). Consistently, IL-6 expression showed no significant change throughout the gut, which is consistent with its expression pattern in bone and serum (Fig. 5 a, b, c and d). Taken together, these data demonstrated that the use of *P.histicola* is related with the expression changes of the osteoclastogenic cytokines site-specifically in gut which may further regulate osteoclast activity through the GM-bone axis.

**Figure 5.**
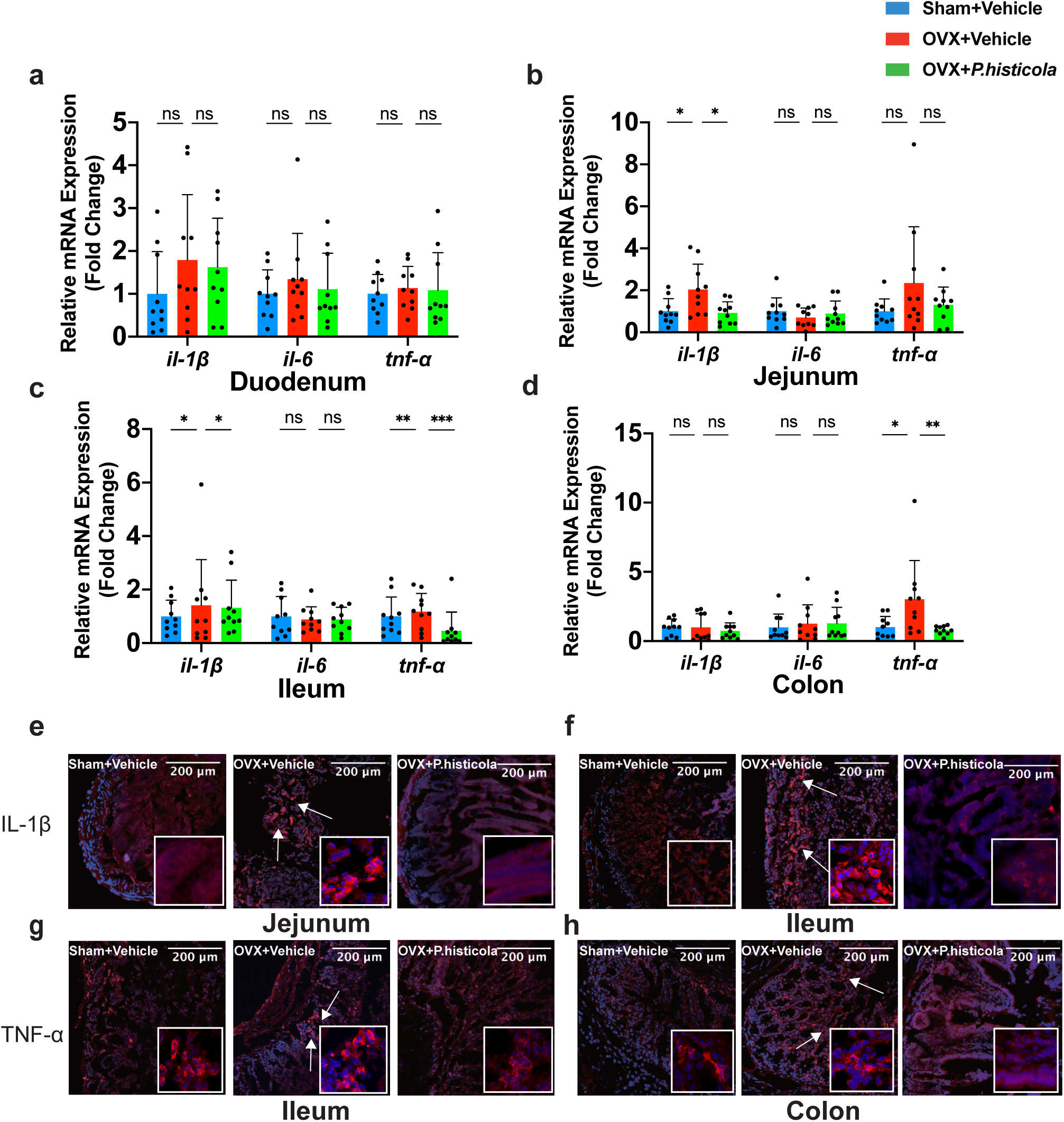
*Prevotella histicola* (*P.histicola*) alters the expression of osteoclastogenic cytokines including IL-1β, IL-6 and TNFα in gut site-specifically. **a** Duodenum, the expression of three osteoclastogenic cytokines didn’t differ among the groups. **b** Jejunum, the IL-1β RNA levels displayed a trend to increase with OVX and a decrease in OVX mice treated with *P.histicola.* **c** Ileum, the IL-1β and TNFα RNA levels both increased with OVX and it could be prevented by *P.histicola* treatment. **d** Colon, the transcript levels of TNFα increased with OVX and *P.histicola* treatment could prevent it. **e-f** Representative immunofluorescence staining images for osteoclastogenesis related inflammatory cytokines IL-1β in mice jejunum and ileum (n=3 per group). **g-h** Representative immunofluorescence staining images for osteoclastogenesis related inflammatory cytokines TNFα in mice ileum and colon (n=3 per group). *** p<0.001, ** p<0.01, * p<0.05. IL-1β, interluekin-1β; IL-6, interluekin-6; TNFα, tumor necrosis factor α; ns, non-significant; P.histicola, Prevotella histicola.

### *P.histicola* ameliorates gut permeability of OVX mice

To elucidate how *P.histicola* prevents OVX induced bone loss via GM-bone axis, we next determined the changes of gut permeability which had been linked to diseases such as IBD and sex steroid deficiency condition (Irwin, Lee et al., 2013, Li, Chassaing et al., 2016a). FITC-labeled dextran was orally given to the mice and its serum level was then detected to show the gut permeability. It was found that the OVX mice has a higher gut permeability compared to the sham group (Fig. 6e), suggesting that estrogen-deficiency may increase gut permeability, which was consistent with previous reports (Li et al., 2016a). *P.histicola* could well maintain the gut permeability as noticed by a similar level of serum dextran with the sham group (Fig. 6e). ZO-1 and occludin are the key proteins which maintain the tight-junction (TJ) integrity of gut(Brun, Castagliuolo et al., 2007). The expressions of ZO-1 and occludin in colon decreased in OVX mice group as showed by the immunofluorescence and WB, which may thus lead to the high gut permeability. However, OVX mice treated with *P.histicola* showed an increase in the expression of TJ proteins ZO-1 and occludin (Fig. 6a-d). Therefore, *P.histicola* prevents OVX induced osteoporosis by ameliorating gut permeability.

**Figure 6.**
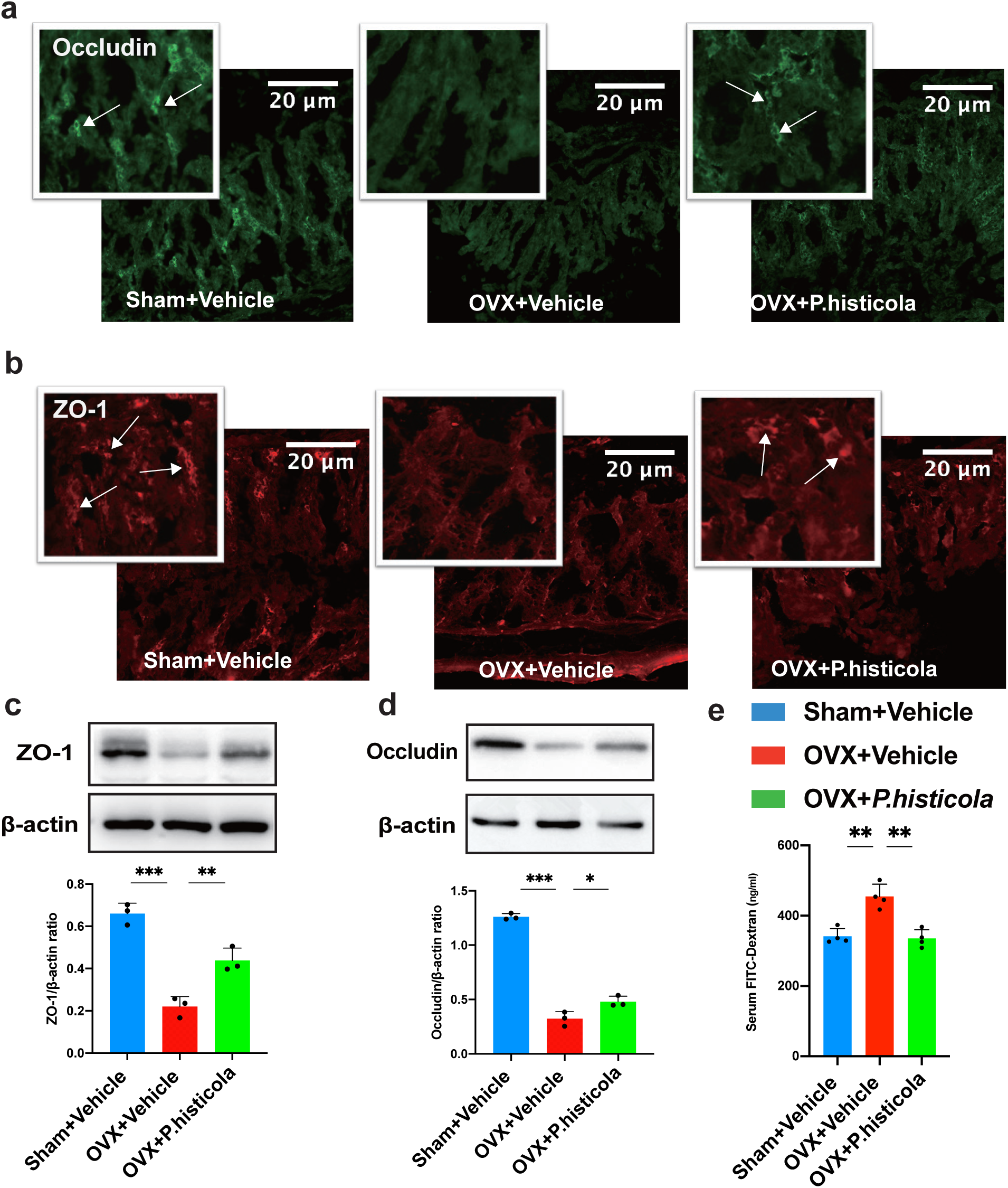
*Prevotella histicola* (*P.histicola*) ameliorates gut permeability of OVX mice. **a-b** Representative immunofluorescence staining images showing the expression patterns of gut tight-junction proteins including occludin and ZO-1 in mice colon (n=3 per group). **c-d** Representative Western Blot images and quantifications of the band intensities showing the changes of ZO-1 and Occludin (n=3 per group). **e** Concentrations of serum FITC labeled dextran in different groups demonstrating the whole intestine permeability (n=4 per group). *** p<0.001, * p<0.05. ZO-1, Zonula Occludens-1; P.histicola, Prevotella histicola; FITC, fluorescein isothiocyanate.

**Figure 7.**
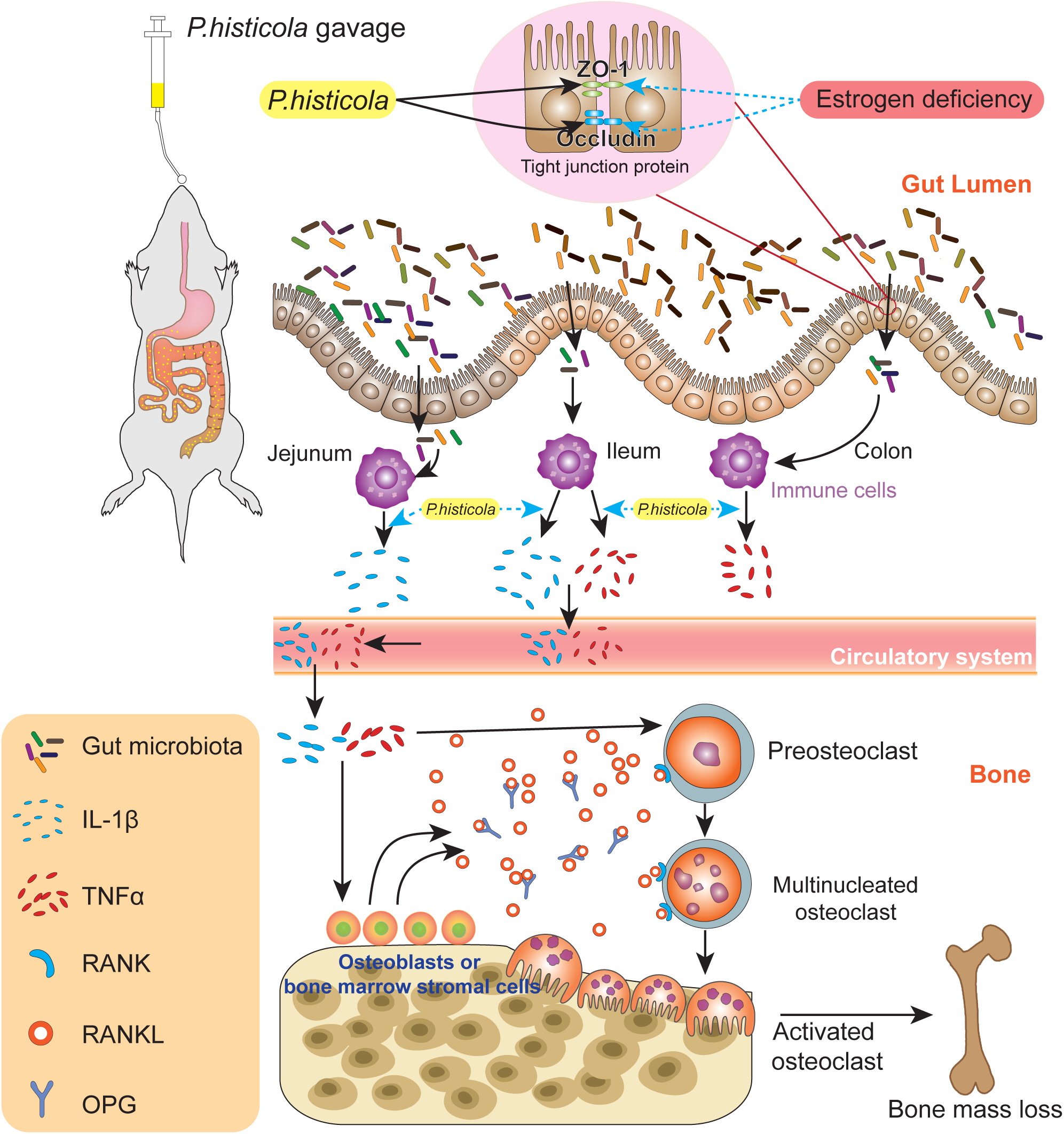
A proposed scheme for the mechanisms of *P.histicola* on preventing estrogen-deficiency induced bone loss via a gut microbiota (GM)-bone axis. *P. histicola* can reduce the intestinal permeability of estrogen-deficient mice by up-regulating the expression of tight junction proteins including ZO-1 and occludin, which further prevents inflammatory cytokines from relasing into circulation. These changes collectively suppressed RANKL-induced osteoclastogenesis thus prevented estrogen-deficiency induced bone loss. IL-1β, interluekin-1β; TNFα, tumor necrosis factor α. RANK, receptor activator of nuclear factor κB; RANKL, receptor activator of nuclear factor κB ligand; OPG, osteoprotegerin; ZO-1, Zonula Occludens-1; P.histicola, Prevotella histicola.

## Discussion

GM is related with a variety of diseases such as diabetes, inflammatory bowel disease and rheumatoid arthritis (Alkanani, Hara et al., 2015, Kostic, Xavier et al., 2014, Scher, Sczesnak et al., 2013). Accumulating studies also found the important role of GM in bone homeostasis (Guss, Horsfield et al., 2017, Ohlsson et al., 2014, Sjogren et al., 2012). Our study firstly investigated and compared the GM richness and diversity between PMO and PMN populations. The results indicated that these two groups shared the similar GM diversity; however, GM richness was significantly altered in PMO patients, suggesting there are some associations between GM and bone. Indeed, it was demonstrated that estrogen deficiency could undermine the bone properties through GM-bone axis (Flores, Shi et al., 2012, Fuhrman, Feigelson et al., 2014). However, the underlying mechanisms still remain largely unknown. In the present study, we identified that the richness of GM in PMO patients was featured by a remarkable decline of genus *Prevotella*, suggesting it may serve as a type of probiotic and have protective effects on the development of postmenopausal osteoporosis.

Previous studies have shown that many probiotics, such as *Lactobacillus reuteri, Lactobacillus paracasei* and the mixture of *Lactobacillus* strains (Britton, Irwin et al., 2014, Ohlsson et al., 2014), can inhibit osteoporosis in animal models, their effects on postmenopausal women are still in doubt. Interestingly, in our study, we found the strain of *Lactobacillu*s showed little difference between two groups. Thus, the protective role *Lactobacillu*s in human may be challenged. Besides, the proportion of *Lactobacillu*s in GM compositions remain very low in our study. We firstly analyzed the GM compositions of PMO patients, aiming to dig out the clinically relevant bacteria which may exhibit protective effects. We found that the genus of *Prevotella* had a much larger proportion in PMN population compared to that in PMO (Fig. 2f). *Prevotella* was first divided from the genus of *Bacteroides* in 1990 and featured by the sensitivity as well as moderately saccharolytic potential to bile (Shah & Collins, 1990). As a type of commensal bacteria, *Prevotella* widely exist in human gut, oral and reproductive tract. Some species of *Prevotella* are associated with human diseases, in particular *Prevotella copri* which has been demonstrated to lead to an increased risk of rheumatoid arthritis (RA) (Maeda, Kurakawa et al., 2016, Scher et al., 2013). Besides, *P.histicola* was proved to inhibit the development of inflammatory arthritis in humanized mice by modulating the gut and system immune response and lowering the gut permeability (Marietta et al., 2016). Moreover, it could also suppress central nervous system inflammatory and ameliorate the symptoms of demyelinating disease (Mangalam et al., 2017). High gut permeability was thought to cause the increased gut inflammatory and subsequently affect bone metabolism (Li et al., 2016a, Schepper, Collins et al., 2019). Therefore, we hypothesized that *P.histicola* may serve as a protective agent on estrogen-deficiency induced bone loss through the GM-bone axis.

To address our hypothesis, *P.histicola* was orally given to the mice following OVX which mimic the postmenopausal estrogen-deficiency. Our data indicated that OVX procedure led to an increased gut permeability and systemic inflammatory-like response in mice, and *P.histicola* treatment could effectively reverse these changes and significantly prevent bone loss via suppressing the osteoclast activity and the expression of osteoclastogenic cytokines. The establishment of osteoporosis animal model is evidenced by the significant changes in parameters of bone microstructure such as Tb.Th, Tb.N, BV/TV, and Tb.Sp (Abdul-Majeed, Mohamed et al., 2015, Muhammad, Luke et al., 2012). The supplement of *P.histicola* could ameliorate these changes and prevent OVX induced bone loss. Excessive osteoclast activity is thought to be the major contributor to the estrogen-deficiency induced bone loss^3^. Bone histomorphometry analysis showed that *P.histicola* reduced the osteoclast number and area on the bone surface, which indicated that *P.histicola* may prevent bone loss by targeting osteoclasts. To better understand the changes of bone remodeling following *P.histicola* treatment, we further analyzed related serum biomarkers. Serum CTX-1 is produced by osteoclasts during the bone resorption process and is a bone resorption marker (Szulc & Delmas, 2008); and P1NP is an osteoblast-derived protein which can reflect the activity of new bone formation (Krege, Lane et al., 2014). In the present study, we found that *P.histicola* could decrease the CTX-1 levels in the serum but with little effect on the P1NP levels. Therefore, we found that the oral administration of *P.histicola* may optimize the GM compositions which subsequently inhibited hyperactive osteoclasts in OVX mice.

Further mechanistic studies revealed that the expression of some indispensable factors for osteoclastogenesis such as RANK and the ratio of RANKL/OPG were decreased due to the treatment of *P.histicola*. RANKL is the major contributor to the differentiation and fusion of osteoclast progenitors by binding to RANK. OPG acts as a soluble decoy receptor majorly derived from osteoblasts and bone marrow stromal cells. OPG competes with RANK in binding to RANKL and prevents osteoclastogenic effect (Boyle et al., 2003). These findings indicated that *P.histicola* decreased the activity of osteoclasts via modulating the RANK/RANKL/OPG pathway. Previous studies have reported that the expression of RANK, RANKL and OPG were affected by some inflammatory cytokines, also termed as osteoclastogenic cytokines, including TNFα, IL-1β and IL-6 (Hofbauer et al., 1999, Kwan Tat et al., 2004, Pacifici, 2016, Sherman et al., 1990). Based on the close relationships between GM and immune system (D’Amelio & Sassi, 2018), we hypothesized that the changes of RANK/RANKL/OPG pathway in the bone were partly regulated by osteoclastogenic cytokines. Next, we assessed the concentrations of serum osteoclastogenic cytokines and the results showed that *P.histicola* could suppress the levels of TNFα and IL-1β in OVX mice. Taken together, these findings indicated that *P.histicola* could reduce the systemic levels of osteoclastogenic cytokines and then inhibited the expression of bone pro-resorptive cytokines, eventually resulting in reduced osteoclast-mediated bone resorption.

How did *P.histicola* suppress the systemic inflammatory response? Previous studies have shown that estrogen deficiency can damage the TJs between epithelial cells, leading to an increased gut permeability. TJ proteins are the fundamental paracellular pathway structure and their integrity is essential to maintain a normal intestinal permeability (Ulluwishewa, Anderson et al., 2011). Increased gut permeability causes an invasion of gut lumen antigenic components and subsequent inflammatory responses (Xu, Jia et al., 2017). The active immune cells lining at the gut lumen release some soluble osteoclastogenic cytokines into the circulation and modulate systemic bone metabolism (Li, Toraldo et al., 2007, Pacifici, 2016). In the present study, we found that the treatment of *P.histicola* enhanced the expression of TJ proteins in colon after OVX, which accounted for a decreased gut permeability. Interestingly, *P.histicola* also reduced the expression of osteoclastogenic cytokines in the gut site-specifically, with IL-1β being majorly affected in jejunum and ileum, and TNFα in the ileum colon. In consistency, the expression of TNFα in the bone was increased in OVX mice and could be inhibited by *P.histicola* treatment. It remains to be further investigated how these cytokines were released and regulated. Nevertheless, it was previously reported that gut activated immune cells could migrate to bone tissues and release TNF*α* or other osteoclastogenic cytokines to regulate bone remodeling (Xu et al., 2017). These findings indicated that *P.histicola* could strengthen intestinal barrier and reduce intestinal inflammation by refraining from the gut-originated osteoclastogenic cytokines into the circulation.

In summary, our study identified for the first time that the significant loss of genera *Prevotella* in postmenopausal osteoporosis patients, suggesting its protective role on osteoporosis. *P.histicola*, a typical specie of *Prevotella*, was then demonstrated to significantly prevent OVX induced bone loss by ameliorating osteoclastic bone resorption through GM-bone axis. Thus, *P.histicola* may serve as a novel probiotic and exhibit a therapeutic effect for osteoclast-related bone disorders.

## Materials and Methods

### Participants recruitment and bone mineral density (BMD) detection

Participants are recruited from The First Affiliated Hospital of Wenzhou Medical University and they were all living in Wenzhou city with similar diet habits. Dual X-ray absorptiometry (DXA) was performed to detect the BMD of lumbar vertebrae and femoral neck. All the subjects are between 55 and 65 years old. The patients with any malignancy, diabetes, astriction, kidney disease and any prebiotics, probiotics or antibiotic treatment within three months were excluded. Forty-two patients were finally included in our study (n=24 in PMO group; n=18 in PMN group) (Table 1). The study was approved by The First Affiliated Hospital of Wenzhou Medical University, Clinical Research Ethics Committee. Informed content was obtained from each participant in this study.

### Sample collection and 16S rRNA gene sequencing

Feces were collected in sterile paper boxes, about 2g samples were transferred to sterile plastic tubes and stored at -80°C immediately. DNA was extracted by QIAamp DNA Stool Mini Kit (QIAGEN, Hilden, Germany)according to the manufacturer’s instructions. The bacterial genomic DNA was used as the template to amplify the V4–V5 hypervariable region of the 16S rRNA gene with the forward primer (515F 5’-GTGCCAGCMGCCGCGGTAA-3’) and (926R 5’-CCGTCAATTCMTTTGAGTTT-3’). Each sample was independently amplified three times. Finally, the PCR products were checked by agarose gel electrophoresis, and the PCR products from the same sample were pooled. The pooled PCR product was used as a template, and the index PCR was performed by using index primers for adding the Illumina index to the library. The amplification products were checked using gel electrophoresis and were purified using the Agencourt AMPure XP Kit (Beckman Coulter, CA, USA). The purified products were the indexed in the 16S V4–V5 library. The library quality was assessed on the <u>Qubit@2.0</u> Fluorometer (Thermo Scientific) and Agilent Bioanalyzer 2100 systems. Finally, the pooled library was sequenced on an Illumina MiSeq 250 Sequencer for generating 2×250 bp paired-end reads.

### Bioinformatics and statistical analysis

The raw reads were quality filtered and merged with the following criteria: (1) Truncation of the raw reads at any site with an average quality score < 20, removal of reads contaminated by adapter and further removal of reads having less than 100 bp by TrimGalore, (2) The paired end reads are merged to tags by Fast Length Adjustment of Short reads (FLASH, v1.2.11), (3) Removal of reads with ambiguous base (N base) and reads with more than 6 bp of homopolymer by Mothur, (4) Removal of reads with low complexity to obtain clean reads for further bioinformatics analysis. The remaining unique reads were chimera checked compared with the gold.fa database (http://drive5.com/uchime/gold.fa) and clustered into operational taxonomic units (OTUs) by UPARSE with 97% similarity cutoff. All OTUs were classified based on Ribosomal Database Project (RDP) Release9 201203 by Mothur. Rarefaction analysis and alpha diversities (including Shannon, Simpson and InvSimpson index) were analyzed by Mothur. Sample tree cluster by Bray-Curtis distance matrix and unweighted pair-group method with arithmetic means (UPGMA) and Jaccard principal coordinate analysis (PCoA) based on OTUs were performed by R Project (Vegan package, V3.3.1). Redundancy analysis (RDA) was analyzed by Canoco for Windows 4.5 (Microcomputer Power, NY, USA), which was assessed by MCPP with 499 random permutations.

### Animal model and experimental design

All the animal experiments were approved by the Animal Experimental Ethical Inspection of Laboratory Animal Centre, Wenzhou Medical University. 10-week-old female C57BL6/J mice were maintained under specific pathogen free (SPF) conditions, with a strict 12h light/dark cycle. The mice had free access to food and water. Mice were allowed to acclimatize to the animal facility for 1 week prior to the start of the OVX surgery. All mice were randomly divided into three groups: Sham procedure +Vehicle (n=10), OVX+Vehicle (n=10) and OVX+*P.histicola* (n=10). The sham ovariectomy is just exteriorized the ovaries but not resect them. After the surgery, the mice had 1 week to recover.

### *P.histicola* culture and supplementation

*P.histicola* dsm 19854 from DSMZ (Deutsche Sammlung von Mikroorganismen und Zellkulturen) was cultured under anaerobic condition in PYG medium (modified) at 37°C for a maximum of 24 hours. For the general storage, the bacteria were mixed with the same volume of sterile glycerin to avoid frozen injury and kept in -20°C. For gavaging, one bacteria tube will be cultured in 10ml medium at 37°C for 20 to 24 hours until the *P.histicola* density was up to 1×10^10^ CFU/ml (OD600=0.723). The mice in OVX+*P.histicola* group were orally gavaged 0.1ml of bacteria(1×10^9^ CFU)every other day throughout the study (12 weeks), while Sham+Vehicle and OVX+Vehicle groups received culture medium only.

### Serum collection and analysis

At the end of experiment, the mice were anesthetized and the blood were taken by heart puncture, the blood was clotted at 37°C for 10 min, put the blood into 4°C for 24 hours and centrifuged at 6000 rpm for 10 minutes to separate the serum and cells, serum was obtained and frozen in -80°C for further analysis. The serum analyses were performed using ELISA kits (Westang, Shanghai, China) according to the manufacturer’s instruction. Cytokines including TNF*α*, IL-1β, IL-6, bone turnover markers CTX-1, and P1NP were analysed in this study.

### Micro-CT scanning and analysis

The mice femurs were collected, fixed and scanned using a Skyscan 1176 micro-CT instrument (Bruker microCT, Kontich, Belgium) as we previously described (Chen, Qiu et al., 2019). The images were then reconstructed with NRecon software (Bruker microCT, Kontich, Belgium). The volume of interest (VOI) was generated between 0.5 ∼1.5 mm above the growth plate of the distal femur. The trabecular bone region of interest (ROI) within this volume was defined and bone parameters including bone volume per tissue volume (BV/TV), trabecular number (Tb.N), trabecular thickness (Tb.Th), and trabecular separation (Tb.Sp) were analyzed by the program CTAn (Bruker microCT, Kontich, Belgium).

### Bone histomorphometry analysis

Following micro-CT analysis, femurs were decalcified in 14% EDTA (Sigma-Aldrich, Sydney, NSW, Australia) at 37 °C for 7 days, and then transferred to an automated vacuum tissue processor (TP-1020, Leica Microsystems, Germany). Sample were then embedded into paraffin for sectioning. Tartrate-resistant acid phosphatase (TRAP) activity staining were performed according to the manufacturer’s protocol (Solarbio, Beijing, China). Section images were acquired using Olympus BX51 (Olympus Corporation,Takachiho,Japan). TRAP-positive osteoclast number (Oc.N/BS) and surface (Oc.S/BS) were quantitated by ImageJ (Rawak Software Inc., Stuttgart, Germany). The skulls of mice were removed from the bodies and removed all the soft tissue, washed them by phosphate buffer saline for 3 times. Fixed with 10% neutral buffered formalin for 24 hours, washed with PBS for 3 times. The skulls were incubated in TRAP activity staining reagent at 37°C for 30 minutes and washed with tap water. Pictures were taken by Sony rx100m3 (Sony, Tokyo, Japan). For the undecalcified sections, we performed in accordance with a previous study (Gao, Zhang et al., 2016). Femurs were proceeded through a serial of gradient ethanol and cleared with toluene, followed by the embedding in methyl methacrylate (MMA) to allow polymerization. Representative sections were vertically prepared by a diamond saw with a thickness of 5 mm and slides were stained with Van Gieson solution containing 1.2% trinitrophenol and 1% acid fuchsin.

### Quantitative real-time PCR (qRT-PCR)

The proximal 1cm of the duodenum, jejunum, ileum, colon, right tibia were used for RNA extraction. The samples were removed from the mice and stored in -80°C. Total RNA was isolated from the tissues using Trizol reagent (Life Technologies, Carlsbad, CA, America). The RNA quality was assessed by detecting the absorbance ratio at A260/230 and A260/280 read between 1.8 and 2.0. RNA was reverse transcribed to cDNA by PrimeScript RT Master Mix (TaKaRa, Beijing, China) according to the manufacturer’s protocol. QRT-PCR reaction in 20ul reactions containing 10ul TB Green Premix Ex Taq II(Tli RNaseH Plus)(TaKaRa, made in china), 3µl of cDNA and 0.4uM forward primers and 0.4uM reverse primers. A melting curve was performed for each primer pair to identify the specific of the primers. Assays were performed in analytical triplicates using a QuantStudio 6 Flex Real-Time PCR System (Applied Biosystems, Warrington, Cheshire, UK) and target gene expression levels were normalized to average expression of housekeeping gene GAPDH using ΔΔCT method. The primer sequences were listed in Table 2.

**Table 2.**
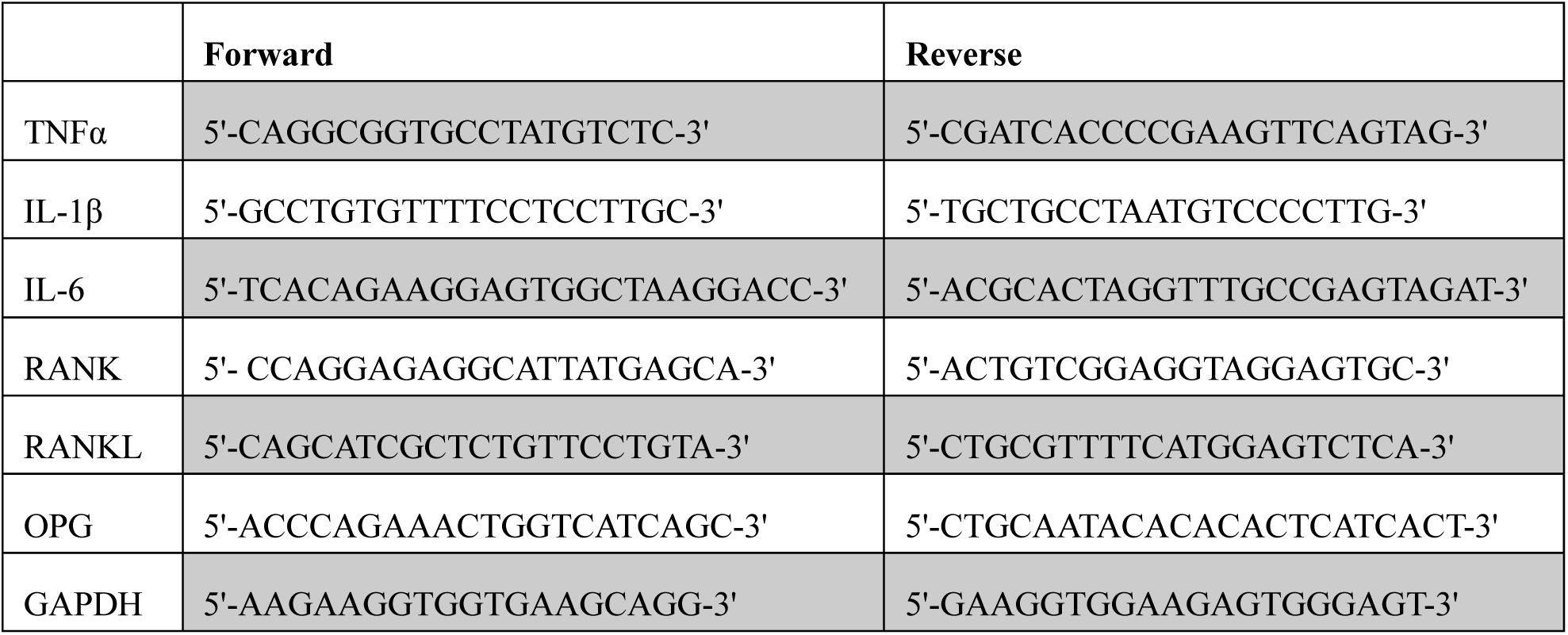
Primers sequences

### Gut permeability analysis

Mice were gavaged FITC-labeled dextran (Sigma-Aldrich, Missouri, USA) (0.6 mg /g) 3 hours before serum collection. The dextran concentration of serum was detected by a fluorescence spectrophotometer (Thermo Scientific Varioskan LUX, Massachusetts, USA) at an excitation wavelength of 485nm and emission at 530nm.

### Western blot

Tissue were lysed by ice-cold RIPA Buffer (Beyotime, Shanghai, China) after weighing and incubated at 4°C for 30 minutes. Following centrifugation at 12 000 × g for 10 minutes, the supernatant was collected and mixed with 5× loading buffer. The proteins were heated at 100°C for 8-10 minutes for denaturation. Next, proteins were separated using 10% sodium dodecyl sulphate-polyacrylamide gel electrophoresis (SDS-PAGE) and transferred onto nitrocellulose membranes (Pallcorporation, New York, USA). After blocking in TBST containing 5% skim milk for 1 hour, the immunoblots were incubated with different primary antibodies including anti-occludin (ab216327, 1:1000, Abcam, USA) or anti-ZO-1 (ab96587, 1:500, Abcam, USA) at 4°C overnight. Subsequently, the membranes were washed three times in TBST, and incubated with the horseradish peroxidase (HRP)-conjugated secondary antibodies for 1 hour. After washing in TBST for another three times, the protein signals were detected using the ECL detection kit (Bio-Rad, California, USA). Blots were analyzed using Quantity One software (Bio-Rad, California, USA).

### Gut immunofluorescence analysis

The proximal 1cm of jejunum, ileum and colon were removed from the body, washed with PBS, fixed with 10% neutral buffered formalin for 24 hours. The samples were embedded and frozen in O.C.T. compound (SAKURA, USA). 10-µm-thickness sections were generated and air-dried at room temperature before staining. For gut tight junction proteins staining, the sections of colon were incubated with rabbit anti-occludin (ab96587, 1:200, Abcam, USA) or rabbit anti-ZO-1 (ab216327, 1:200, Abcam, USA) for 2 h and probed with Alexa Fluor 488 AffiniPure Donkey Anti-Rabbit antibodies (1:500, Jackson, USA) or Alexa Fluor 594 AffiniPure Donkey Anti-Rabbit antibodies (1:500, Jackson). For gut osteoclastogenic cytokines staining, the jejunum and colon sections were incubated with Mouse Anti-IL-1β or Mouse Anti-TNFα (sc-52012, sc-52746, 1: 200, Santa Cruz, USA), the ileum sections were divided into two groups, one group with Mouse Anti-IL-1β and another with Mouse Anti-TNFα. All sections were incubated overnight at 4 °C prior to the incubation with Alexa Fluor 594 AffiniPure Donkey Anti-Rabbit Antibodies (1:500, Jackson, USA) or Alexa Fluor 594-AffiniPure Goat Anti-Mouse Antibodies (1: 500, Jackson, USA). Sections were observed and pictured by a fluorescence microscope (Olympus Corporation,Takachiho,Japan).

### Statistical Analysis

All statistical analyses were performed with SPSS 22.0 for Windows (SPSS Inc., USA). In the GM diversity analysis, we used an independent-sample t test and the Mann–Whitney test. Correlations were determined with Spearman’s correlation. The resulting p values were adjusted using the Benjamini–Hochberg false discovery rate (FDR) correction. Only FDR-corrected p values below 0.05 were considered significant. In the animal experiment, quantitative data were presented as mean ± SD. Statistical significance was determined by independent-sample t test. A p-value of less than 0.05 was considered to be significant.

## Acknowledgements

This study was supported by the Zhejiang Provincial Natural Science Foundation of China (LGF19H070004) and Medical Health Science and Technology Project of Zhejiang Provincial of China (2019KY446). We acknowledge the facilities and technical assistance of the Centre for Microscopy, Characterization & Analysis, the University of Western Australia.

## Author contributions

L.C., K.J. conceived and designed the experiments. Z.W., C.W. analyzed the fecal samples. Z.W., K.C., J.C., H.P. and J.D. performed the animal study. Y.L, P.W, J.Y. and J.L. analyzed the data. Z.W. and K.C. prepared the data and figures with the help of all authors. Z.W. and K.C. drafted the manuscript, and L.C., K.J., J.X revised the manuscript. L.C., K.J., J.X supervised and coordinated the project.

## Data availability

The authors declare that all the data supporting the findings of this manuscript are available within the paper and its supplementary information. 16S RNA sequencing data are deposited in publicly accessible database and available under the following accession code: PRJNA631117.

